# Clustering Compresses Attractors in Watts-Strogatz Threshold Boolean Networks

**DOI:** 10.64898/2026.01.01.697270

**Authors:** Maram Alqarni, Mark Cooper, Diane Donovan, James Lefevre

## Abstract

**Abstract:** Does higher clustering shorten attractor periods? We examine whether the global clustering coefficient *C*, a direct measure of triangle density, predicts attractor lengths in synchronous, signed-threshold Boolean networks on Watts-Strogatz (WS) graphs. We generate 330 directed, signed WS networks spanning sizes *N* = 10–100 and mean degrees 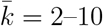, simulate dynamics from *M* = 100 random initial states per graph with exact cycle detection, and summarise each graph by the average log attractor period (equivalently, the geometric mean period). Our primary analysis relates this log-period summary to *C* while adjusting for *N*, 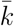, the mean directed shortest path (MSP), and including nonlinear size-degree and clustering-degree interactions.

**Main finding:** Higher clustering robustly shortens attractor periods: a 0.10 increase in *C* (*C* ∈ [0, 1]) corresponds to an ≈ 13–15% lower expected geometric mean period, and moving from *C* = 0.000 to *C* = 0.460 yields an ≈ 50% reduction, holding other properties fixed. The effect persists when the linear *C* term is replaced by a nonlinear function of *C*, and it replicates in held-out graph instances (graphs not used to fit the model). Shorter periods are not explained by an increase in fixed points under the strict comparator (*>*); rather, higher triangle density shifts mass from long cycles to medium-length cycles.

**Significance:** In threshold-like logic, settling speed and oscillatory stability are central to computation and control. Our results provide a direct, quantitative link between triangle density and these long-run behaviours, showing that *C* acts as a structural lever on temporal complexity.

*All figures and tables are reproducible from the accompanying code, data, and analysis scripts*.

## 1 Introduction

## 1.1 Boolean networks and attractors

Boolean networks (BNs) provide a compact language for studying how the wiring of a regulatory system shapes its long-term behaviour [1, 2, 3]. Each node holds a binary state (on/off); at discrete time steps all nodes update according to local logical rules that encode activation and repression. Despite their simplicity, such models capture essential features of biological regulation when kinetic parameters are unknown or noisy, and they offer transparent links between structure and dynamics through the lens of attractors-fixed points and periodic cycles that represent stable phenotypes or recurring activity patterns [4, 5]. Kauffman’s seminal work introduced random Boolean networks as abstract models of genetic regulation, proposing that stable cellular phenotypes correspond to attractors in these simplified networks [3]. Subsequent studies formalized the attractor concept and showed that network connectivity and logic determine whether dynamics settle into an ordered model (with many fixed points and short cycles) or a chaotic model (with few fixed points and long, complex cycles) [2, 6, 7, 8, 9, 10]. Biological networks are thought to operate near this critical balance between order and chaos, ensuring both stability and adaptability, and indeed BNs have been applied to real systems to capture such dynamics. For example, a Boolean model of the yeast cell cycle network reproduces its robust oscillatory progression through cell cycle states [10, 11].

## 1.2 Clustering and small-world structure

One structural feature that may modulate a network’s dynamical model is clustering. Clustering measures the prevalence of closed triangles in a network, the tendency for neighbours of a node to also be connected with one another and reflects the density of local feedback cycles [12, 13]. Many natural networks, including biological regulatory networks, exhibit significant clustering [14, 15, 16]. In small-world networks, introduced by Watts and Strogatz (1998), high clustering combines with short path lengths, enabling rapid communication while maintaining local community connection [17]. Because clustering directly counts local feedback motifs (triangles) and can be tuned in the WS models, it is a natural structural axis on which to study attractor statistics [17, 18].

### 1.3 Prior work and mechanistic expectations

Prior work has linked small-world structure to ordered dynamics, but mostly through indirect measures. However, most studies evaluate proxies (storage, damage) rather than directly quantifying how the clustering coefficient *C* relates to attractor period length. For these reasons, we hypothesise that increasing clustering should shift a Boolean network’s dynamics toward the ordered side, yielding shorter attractor cycles on average.

### 1.4 Modelling framework: threshold dynamics on directed WS graphs

We address this gap by testing the clustering–period link using a biologically grounded update rule. Threshold logic is widely used to approximate regulatory interactions because it mirrors the sigmoidal responses and saturations common in gene regulation and signalling [19, 5, 20]. Majority-like threshold rules reduce effective sensitivity and often promote reliable settling, a tendency also observed in networks enriched for canalising functions, where a single input can determine the output regardless of others [21, 9, 22]. In fact, canalizing rules, which stabilize dynamics and prevent chaos, are frequently included in carefully constructed biological Boolean models [5, 20]. Recent theory on threshold Boolean networks further clarifies stability conditions and supports using these rules to probe structure dynamics links [23, 24]. For these reasons, we adopt synchronous signed-threshold updates and vary network structure while keeping the rule class fixed. This design isolates the effect of clustering without confounding it with changes in logical update properties.

Our setting is the directed WS model, analysed on the scale of the global clustering coefficient *C* (see Sec. 4). The same *p* can produce different triangle densities across sizes and degrees, whereas *C* is directly comparable between instances [17, 15]. Moreover, empirical networks do not come with a rewiring knob but do have measurable clustering [14, 16]. In this study we estimate how the log–mean period 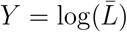 varies with *C* while adjusting for network size *N*, average degree 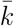, and mean directed shortest path (MSP); we also examine average betweenness centrality (ABC) in exploratory variants to enable fair comparisons across mixed geometries [15, 8]. Estimation details and diagnostics are provided in Sec. 4, and substantive findings are presented in Sec. 5.

### 1.5 Contributions

This paper makes four contributions. (1) *Direct measurement of clustering–period links:* we quantify how *C* relates to the attractor period *L* under synchronous signed-threshold dynamics on directed WS graphs, moving beyond proxies such as AIS and damage spreading [25, 26]. (2) *Effect sizes and generalisation:* we report interpretable ratios on the log period scale and assess out of sample generalisation across graph instances [27]. (3) *Distributional consequences:* period shortening arises primarily from a shift of probability mass away from long cycles toward medium length cycles; critically, with jittered thresholds ties are almost surely absent (so *>* and ≥ yield effectively identical dynamics), and the frequency of fixed points does not increase under the strict comparator (*>*), clarifying how shorter periods emerge in practice [2, 28, 4]. (4) *Reproducible detection:* we provide exact attractor detection via a hash-and-backtrack scheme, with parameters and stopping rules specified for replication. The method stores each visited state in a hash table and, when a repeat is found, backtracks to the first occurrence, returning the *exact* cycle and its period. Common alternatives include SAT/BDD-based exact enumerators and state-space sampling (approximate), which differ mainly in memory use and scalability [4, 29].

## 2 Background

This section outlines the conceptual background for how clustering shapes attractor periods in threshold Boolean networks and explains why we focus on directed, signed Watts–Strogatz (WS) graphs.

### 2.1 Boolean networks, attractors, and phases

A classic result is that random Boolean networks exhibit distinct dynamical phases depending on their parameters. In an ordered phase, perturbations to the state of one node tend to die out over time; such networks have many fixed points and generally short cycles. In a chaotic phase, small perturbations amplify and propagate indefinitely, resulting in long transients and complex cycles [2]. Separating these extremes is a critical edge-of-chaos model where the network is maximally sensitive yet not fully chaotic [10, 30, 31]. Kauffman [3] first noted that random networks with low connectivity often settle quickly into point attractors, whereas highly connected networks can exhibit chaotic, lengthy oscillations. Later work formalised this via the notion of average sensitivity of the Boolean update rules; if the average sensitivity is less than one (for example, due to canalisation or bias towards stable outputs), the network is ordered; if it is greater than one, the network is chaotic [9]. Shmulevich and Kauffman [9] quantitatively linked sensitivity to Boolean logic types, and Klemm and Bornholdt [7] observed that networks can have both stable and unstable attractors depending on structure and bias. These insights explain why real genetic networks are believed to operate near an ordered model [11]. Where relevant, we use signed–threshold dynamics to ground later comparisons in a biologically common rule class.

### 2.2 Small-world structure, clustering, and the WS baseline

Beyond node-level logic, the topology of interactions influences a Boolean network’s dynamics. Many empirical networks are neither completely random nor completely regular, but instead have small-world structure combining high clustering with short path lengths [17, 15, 25, 26]. The Watts–Strogatz (WS) model generates small-world networks by starting from a ring lattice and randomly rewiring a fraction of edges to create shortcuts [17]. At intermediate rewiring, the network retains most local links and thus high clustering while gaining a few long-range links that shrink distances [18].

Our focus on WS complements other network families. In Erdős–Rényi graphs, clustering is low and largely a function of size; mean degree is dominant [15]. In scale-free graphs, degree heterogeneity changes sensitivity and stability; Barabási and Albert showed that hubs shape connectivity [32], and Aldana analysed Boolean dynamics on scale-free topology [6]. Modular organisation can enhance robustness by confining perturbations within modules [33]. To vary triangle density at nearly constant degree and path length, we use WS as the ideal model [17, 18]. In other families, clustering co-varies with other graph properties, making it hard to isolate the effect of clustering alone. Our approach is complementary to motif-level studies predicting attractors from small circuit templates [34]: where motif analyses are local and combinatorial, our study is global and statistical at the network level, asking how clustering relates to typical attractor periods across many graphs.

### 2.3 Mechanistic lenses linking clustering to periods

Although different in methods, three lines of reasoning converge on the prediction that increasing clustering should shorten attractor periods.

#### Information storage and redundancy

In clustered neighbourhoods, a node’s inputs are interconnected, creating overlapping paths of predictive information; small-world structure with high clustering raises active information storage (AIS) [25]. In WS networks, nodes in highly clustered regions have higher AIS, reflecting stronger local memory. Under threshold logic, such redundancy reduces effective sensitivity, supporting shorter cycles.

#### Damage spreading and sensitivity

Damage spreading tracks how an initial perturbation propagates. Ferraz and Herrmann [35] examined Boolean dynamics on small-world graphs and observed that introducing even a small fraction of shortcuts could drive a transition toward more chaotic behaviour as damage spreading increased. Lu and Teuscher [26] demonstrated that adding local links, thereby increasing clustering, suppresses damage spreading, whereas adding long-range links, thereby decreasing clustering, amplifies it. Lower sensitivity implies shorter transients and fewer opportunities to traverse long cycles [2].

#### Loop closure and feedback motifs

Analyses connect the probability of closing onto a short cycle to local feedback and redundancy [36]. Motif-level studies in related threshold linear systems show that small feedback circuits strongly constrain possible attractors [34]. Because clustering counts triangles that form local feedback cycles, higher clustering increases opportunities for early loop closure, shifting probability toward short cycles.

Together, these perspectives offer a coherent mechanistic account consistent with network theory [15]: triangles increase storage and decrease sensitivity; lower sensitivity and more local feedback increase the likelihood of short cycles; therefore, as clustering increases, attractor periods should decrease on average. Our empirical analysis in Sec. 5 tests this prediction directly by relating clustering to attractor periods under a fixed update rule.

### 2.4 Threshold logic, canalisation, and signed edges

In regulatory modelling, Boolean update rules range from unconstrained truth tables to structured, biologically informed families. Threshold rules are widely used in gene regulation and signalling because they capture saturating, sigmoidal responses with few parameters [19, 5, 20]. Majority-type thresholds lower average sensitivity and favour ordered dynamics with reliable settling [5]. Imposing threshold logic yields robust network behaviour even on random topologies [37], and robustness itself has been proposed as an evolutionary organising principle [10]. Canalising functions further stabilise dynamics and are enriched in curated biological Boolean models; networks composed entirely of canalising rules are provably stable [5, 21]. Both threshold logic and canalisation reduce effective sensitivity, tending to shorten transients and limit attractor periods [9].

Many regulatory and signalling networks are naturally represented as directed, signed graphs, with edges encoding activation (positive) or inhibition (negative) [29, 38]. Small-world models admit directed variants in which undirected links are oriented or edges are rewired in a directed manner, typically preserving narrow inand outdegree distributions while retaining short paths and local cohesion [15, 18]. Measures of clustering generalise from undirected transitivity to directed motifs, enabling analyses that distinguish feed-forward triads from directed 3-cycles where appropriate [15, 12]. In Boolean formulations with signed thresholds, edge signs act as positive or negative weights on inputs, offering a simple and biologically interpretable way to model activation and repression; when sign information is unavailable, a common neutral baseline assumes a roughly equal mix of activating and inhibiting edges [5, 37]. These considerations motivate our choice of synchronous signed-threshold dynamics on directed WS graphs (Sec. 4).

### 2.5 Attractor distributions, proxies for order, and structural covariates

Analyses of Boolean dynamics commonly characterise attractor landscapes using *distributional* summaries rather than means alone. Under a specified initial state distribution and update scheme, one considers the probability of fixed points Pr[*L* = 1] and the probability of short nontrivial cycles (e.g., Pr[*L* ≤ 4]), together with full distributional views such as empirical cumulative distribution functions (ECDFs) of *L* [4, 29]. These summaries reveal whether structural changes (e.g., increased clustering) primarily shift mass away from long cycles toward shorter or moderate cycles, complementing aggregate measures such as the mean period and, for modelling purposes, its log transform [4, 27].

Much of the literature assesses dynamical order via *proxies* such as active information storage (AIS) or damage spreading [25, 26]. A complementary line of work measures attractor *periods* directly and treats the period (or its log transform; see *Y* in Eq. 9) as the response, relating variation in *L* to structural features such as clustering while holding other properties in view [15, 8]. Framing results on the period scale translates qualitative expectations about storage and sensitivity into concrete, testable statements about long-run behaviour.

Attractor lengths are influenced by multiple structural features beyond clustering, including network size *N*, average degree 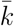, mean (directed) shortest path (MSP; Eq. 3), and average betweenness centrality (ABC; Eq. 5) [15, 8, 39]. Because these quantities often co-vary with clustering under common generative models, empirical analyses typically adjust for them, e.g., modelling log *L* as a function of *C* with *N*, 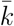, MSP, and ABC as covariates to isolate the partial association of clustering while comparing networks that are similar in scale, sparsity, reachability, and flow centralisation.

## 3 Mathematics Concepts and Definitions

In this section, we introduce the fundamental concepts and definitions that will be used throughout this paper [12, 13, 15, 18, 3, 1, 2, 4].

### 3.1 Graphs, adjacency, and degree

A *graph G* = (*V, E*) has a nonempty *vertex set V* and an *edge set E*. In an undirected graph, *E* ⊆ {{*u, v*}: *u, v* ∈ *V, u* ≠ *v*}; in a directed graph (a *digraph*), *E* ⊆ *V* × *V* with ordered pairs (*u, v*). The *order of G* is *N* = |*V*| and the *size of G* is *m* = |*E*|. Throughout this paper, graphs are simple unless stated otherwise (no self-loops or parallel edges) [12, 13].

An *adjacency matrix* is *A* = [*a*_*ij*_]. For an undirected simple graph, *a*_*ij*_ = *a*_*ji*_ = 1 if {*v*_*i*_, *v*_*j*_} ∈ *E* and 0 otherwise. For a digraph, *a*_*ij*_ = 1 if (*v*_*i*_, *v*_*j*_) ∈ *E* and symmetry is not required. For dynamical models we use a *signed adjacency* with *a*_*ij*_ ∈ {−1, 0, +1} to encode activation (+1) and inhibition (−1).

When measuring *undirected* features (e.g., clustering), we pass to the *underlying undirected simple graph G*^*u*^ with adjacency *U* = [*u*_*ij*_] defined by

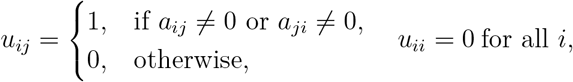

and then treat *G*^*u*^ as a simple graph (so directions and signs are both ignored and parallel edges are not allowed). Unless otherwise noted, any undirected network statistic is computed on *G*^*u*^ constructed from *A* as above [13, 15].

The *degree of vertex v*_*i*_ in an undirected graph is deg(*v*_*i*_) = ∑_*j*_ *u*_*ij*_ and equals the size of the *neighborhood N* (*v*_*i*_) = {*v*_*j*_ : {*v*_*i*_, *v*_*j*_} ∈ *E*}. In a digraph, the *in-degree* and *out-degree* are

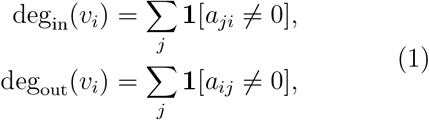

where **1**[·] denotes the indicator, equal to 1 when the condition holds and 0 otherwise. We write the *total degree of vertex v*_*i*_ as deg_tot_(*v*_*i*_) = deg_in_(*v*_*i*_) + deg_out_(*v*_*i*_) (in an undirected graph, deg_tot_(*v*_*i*_) = deg(*v*_*i*_)), and define the *average degree* as 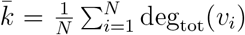 deg_tot_(*v*_*i*_) (so 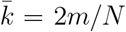 for simple graphs) [12].

### 3.2 Paths, distance, clustering, and path-based measures

A *walk* of *length p* is a sequence (*u*_0_, …, *u*_*p*_) with adjacent consecutive vertices; a *path* is a walk with no repeated vertices [12]. In a digraph, we write *u* ⇒ *v* to denote that there exists a directed path from *u* to *v*. A (simple) *cycle* is a closed walk of length at least 3 whose vertices are all distinct except the first and last. A *3-cycle* (a *triangle*) is a cycle of length 3 (in a digraph, edges of the cycle must respect orientation).

The *distance d*(*u, v*) is the length of a shortest path from *u* to *v* (respecting edge directions in digraphs). The *diameter of G* is the maximum finite distance [15].

For clustering we count *triangles* (3-cycles) and *connected triples* (wedges) on the *underlying undirected simple graph G*^*u*^. A *connected triple (wedge)* is a path of length 2, *u* −*v* −*w*, in *G*^*u*^, with *u* ≠ *w*. Writing *T* for the number of triangles and Λ for the number of connected triples, the global clustering coefficient (transitivity) *C* is defined as:

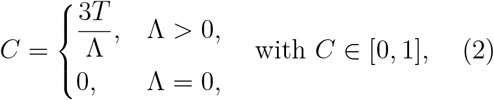

which is the fraction of wedges that are closed into triangles [15, 18, 14].

We use the mean *directed* shortest path (MSP), defined as:

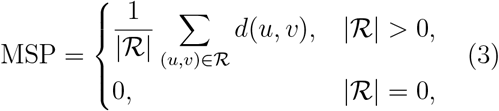

where

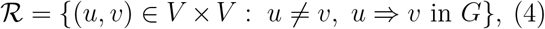

and *d*(*u, v*) is the length of a shortest *directed* path from *u* to *v*. Thus, *G* need not be (strongly) connected; the average is taken only over ordered pairs that are reachable by a directed path (i.e., pairs in ℛ) [15].

The average betweenness centrality is

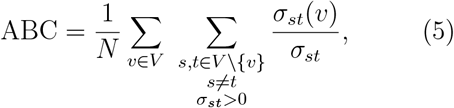

where *σ*_*st*_(*v*) is the number of shortest directed *s* → *t* paths that pass through *v*, and *σ*_*st*_ counts shortest directed *s* → *t* paths from *s* to *t* [40].

### 3.3 Boolean networks and attractors

Given a graph *G* = (*V, E*) with |*V*| = *N*, we associate a Boolean state to each vertex at each discrete time *t* ∈ ℤ_*≥*0_. The *state vector* is

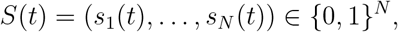

with a chosen initial condition *S*(0) ∈ {0, 1}^*N*^. A Boolean network on *G* is the deterministic map

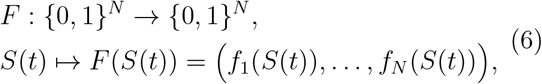

where each coordinate function *f*_*j*_ : {0, 1}^*N*^ → {0, 1} depends only on the states of the in-neighbors of the vertex *v*_*j*_ in *G* [1, 2]. The synchronous evolution is then

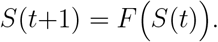

An *update scheme* specifies when node states are updated. In the *synchronous* scheme, all nodes update simultaneously at each integer time step according to Eq. (6). Asynchronous variants (e.g., random-order or general-asynchronous) update subsets or single nodes per step; in this study we focus on synchronous updates [2, 29].

An *update rule* specifies the coordinate functions *f*_*j*_ used in Eq. (6) and, together with the update scheme, determines the next state *S*(*t*+1) from *S*(*t*) [2]. A widely used rule in signed regulatory models is the *signed–threshold rule*,

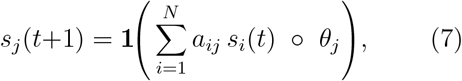

where *a*_*ij*_ ∈ {−1, 0, +1}, *θ*_*j*_ ∈ ℝ is a node-specific threshold, and the comparison symbol ◦ ∈ {*>*, ≥}; in this study we use the strict comparator “*>*” throughout (see Sec. 4) [2, 5, 37].

An *attractor* is a minimal set *A* = {*a*_0_, …, *a*_*L*−1_} ⊆ {0, 1}^*N*^ such that *F* (*a*_*r*_) = *a*_(*r*+1) mod *L*_ for all *r* ∈ {0, …, *L* − 1}. Equivalently, *F*^*L*^(*a*_*r*_) = *a*_*r*_ with no smaller positive period. The *period* is *L* ∈ 1, 2, …} (a fixed point is the case *L* = 1).

The *state transition graph* places a directed edge *S* → *F* (*S*) on the 2^*N*^ states (out-degree 1 per state); attractors are precisely the directed cycles (with fixed points as 1-cycles) [2, 4]. From a given initial state *S*(0), the trajectory *S*(0), *S*(1), … eventually repeats because 0, 1^*N*^ is finite. The *transient length* from *S*(0) is

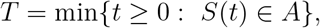

i.e., the first time the trajectory enters the repeating cycle.

When summarizing outcomes over many initial states, we report the *attractor period* and its log transform. From *M* simulations with periods *L*_*r*_,

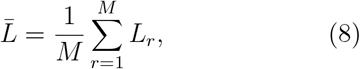

and

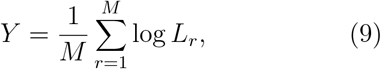

so that the geometric mean attractor period is *L*_geo_ = exp(*Y*) and a difference Δ*Y* corresponds to a multiplicative factor *e*^Δ*Y*^ on *L*_geo_ [4, 27].

### 3.4 Graph models: Watts-Strogatz and Erdős-Rényi

#### Watts-Strogatz (WS) model

Fix an integer *N* ≥ 3 and an integer *κ* ∈ {1, …, ⌊(*N* − 1)/2⌋}, and let *p* ∈ [0, 1]. Let *V* = {0, 1, …, *N* − 1} and for *u, v* ∈ *V* with *u* ≠ *v* define the circular distance

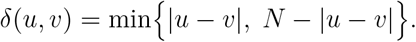

The ring lattice ℒ (*N, κ*) has edge set

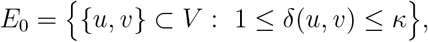

so every vertex has undirected degree deg(*u*) = 2*κ* (i.e., *κ* is the ring-lattice parameter). Construct the undirected WS graph *H* = WS(*N, κ, p*) by independently rewiring each {*u, v*} ∈ *E*_0_ once: with probability *p* replace {*u, v*} by {*u, w*}, where *w* is drawn uniformly from *V \* ({*u*} ∪ *N*_*H*_ (*u*)) at the time of rewiring (avoids self-loops and multiedges); otherwise keep {*u, v*}. This preserves *m* = |*E*(*H*)| = *Nκ* and hence the mean (undirected) degree 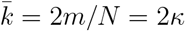 [17, 15, 18].

The *directed* WS graph 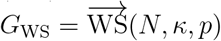 is obtained by orienting each undirected edge of *H* independently and uniformly at random:

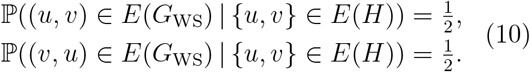

Conditioned on *H*,

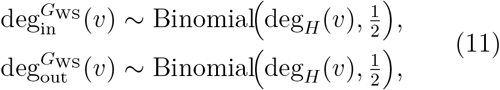

So

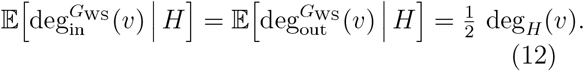

#### Erdős-Rényi (ER) model

Given *N* ≥ 1 and *q* ∈ [0, 1], the undirected model *H* = ER(*N, q*) is defined by independent edges with

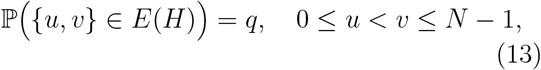

so 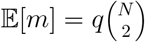 and 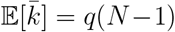 [15, 18]. The *directed* E-R graph 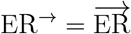 (*N, q*) is obtained by the same orientation rule as in Eq. (10).

#### Matching expected mean degree

For comparisons between WS(*N, κ, p*) and ER(*N, q*), we match expected mean degree by choosing

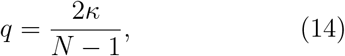

valid whenever 2*κ < N*, so both undirected models have the same expected mean degree 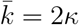 (and thus the same expected total degree after orientation) [15, 18].

## 4 Methods

### 4.1 Graph model: small-world networks and clustering

We use the Watts-Strogatz (WS) small-world model to vary triangle density while keeping the network size *N* and average degree 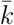 controlled (Sec. 3.4, Def. 3.4; [17, 15, 18]). For comparisons with the baseline we match the expected mean degree using Eq. (14).

To obtain a directed, signed network consistent with regulatory settings, we first generate an undirected WS graph and then orient each edge once using the random orientation rule in Eq. (10), creating exactly one directed arc per undirected edge and no reciprocal *u* ↔ *v* pairs. We then independently assign a sign to each arc, +1 for activation and −1 for inhibition with equal probability, yielding a signed adjacency matrix *A* = [*a*_*ij*_] ∈ −1, 0, +1}^*N ×N*^ [5, 2, 15].

We measure clustering on the underlying undirected simple graph *G*^*u*^ using the global clustering coefficient *C* (Sec. 3, Eq. (2)).

### 4.2 Network generation and sampling

We varied network size over *N* ∈ {10, 20, …, 100}. For each *N*, we considered degree inputs *d* ∈ {1, …, ⌊*N/*10⌋}. Within the WS model (Def. 3.4), we set

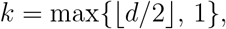

which yields an undirected average degree 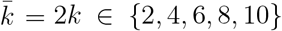. Each (*N, d*) specification was crossed with rewiring probabilities *p* ∈ {0.01, 0.05, 0.10, 0.20, 0.40, 0.60}. The total number of (*N, d*) specifications is

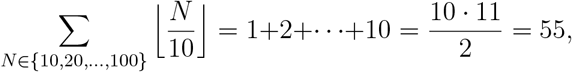

so this design yields 55 × 6 = 330 WS graphs that span topologies from highly clustered small-world networks to nearly random graphs [18].

For baseline comparisons we also generate degree-matched Erdős-Rényi graphs (Def. 3.4) with edge probability *q* chosen via Eq. (14). The main clustering results, however, are reported for the WS ensemble. Across this parameter sweep over *N*, 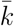, and *p* (with degree-matched ER baselines), the observed clustering on *G*^*u*^ ranges from *C* ≈ 0 (nearly random) to *C* ≈ 0.63 (highly clustered).

### 4.3 Graph properties and controls

For each directed, signed graph, we compute five structural descriptors (Sec. 3): the size *N*; the average degree 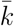; the mean directed shortest path (MSP; Eq. (3)); the average betweenness centrality (ABC; Eq. (5)); and the clustering coefficient *C* (Eq. (2)) evaluated on the underlying undirected simple graph *G*^*u*^. These quantities are defined formally in Sec. 3; here we use them to characterise each graph and as candidate covariates when relating clustering to dynamical outcomes [3, 2, 6, 41, 15, 18, 5, 8, 40]. In the main regression we use *N*, 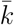, MSP, and *C* as predictors; ABC is computed but only used in exploratory checks and does not enter the primary model. As an initial exploratory step, we also compute Spearman rank correlations between the average log-period *Y* and each structural descriptor (*N*, 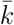, MSP, ABC, *C*) to characterise simple univariate associations and motivate the choice of covariates in the regression models.

### 4.4 Why analyse effects on the *C* scale?

We report effects on the global clustering coefficient *C* rather than the WS rewiring probability *p* because (i) the same *p* can yield different triangle densities across *N* and 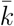, (ii) *C* is directly comparable across sizes and degrees and is measurable in empirical networks, and (iii) statements on the *C* scale transfer beyond a specific generator [17, 15].

### 4.5 Signed-threshold Boolean dynamics and thresholds

We adopt synchronous updates as a standard and tractable baseline for Boolean network analyses and focus exclusively on synchronous dynamics in this study [2, 4]. We use the signed-threshold update defined in Sec. 3, Eq. (7), with a strict “*>*” comparator throughout (the signed adjacency *A* = [*a*_*ij*_] is as defined in Sec. 3).

Node thresholds follow a majority-style rule with a small tie-breaking perturbation:

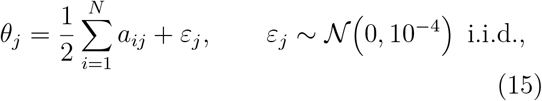

which reduces average sensitivity and promotes stable behaviour in signed regulatory networks [9, 21]. The small perturbation *ε*_*j*_ breaks exact ties without materially changing the dynamics. We keep the strict comparator “*>*” to avoid spurious fixed points at equality [5, 9]. Edge signs are assigned with equal probability to provide a neutral baseline for activation-inhibition balance when system-specific information is unavailable [37, 2]. The update rule is implemented in C++ for efficient simulation and attractor detection.

### 4.6 Simulation and attractor detection

For each graph we run *M* = 100 independent simulations from random initial states *S*(0) ~ Bernoulli(0.5)^*N*^. This choice balances coverage of the state space and computational cost; the Bernoulli(0.5) start is an uninformative prior that matches the sign-balanced wiring [3, 2]. The finite, deterministic state space guarantees eventual periodic behaviour [2].

We detect cycles exactly using a hash-based backtracking method implemented in C++. Each full state vector is stored in a hash table for constant-time lookup; when a state repeats at time *t* with first occurrence at *t*_1_, we report the period *L* = *t* − *t*_1_ after a short backtrack to confirm the earliest repeating state [4]. We impose a conservative cap of 10^8^ update steps per run. If a trajectory has not repeated by the cap, the run *times out* and is excluded from the averages for 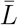 and *Y*; in Eq. (8) *M* is the number of *finished* runs. In our experiments, no run timed out, so all runs were included.

### 4.7 Dynamical outcomes

If run *r* converges to a fixed point, its period is *L*_*r*_ = 1; if it enters an *ℓ*-cycle, its period is *L*_*r*_ = *ℓ >* 1. For each graph we therefore obtain *M* periods *L*_1_, …, *L*_*M*_.

Our primary summary is the *average log-period Y* defined in Eq. (9). Here and throughout, log denotes the natural logarithm (base *e*), and back-transformations use exp(·). With this convention, *L*_geo_ = exp(*Y*) is the geometric mean attractor period for that graph. Differences on the *Y*-scale are multiplicative on *L*_geo_: a change Δ*Y* corresponds to a factor exp(Δ*Y*) on the geometric mean period [27].

For completeness we also define the arithmetic mean period 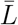 in Eq. (8), but the statistical analyses and figures are based on *Y* and *L*_geo_ unless otherwise noted. In the implementation, *Y* is computed by taking the natural logarithm of each run-specific period *L*_*r*_, averaging over the *M* = 100 runs per graph, and storing the result as `AvgLogPeriod` in the aggregated results file used for the regression. The analysis dataset thus contains one row per graph: the response *Y* and the associated structural descriptors (*N*, 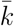, MSP, ABC, *C*). In a small number of descriptive plots of the raw period distribution we use a base-10 logarithmic axis for *L*_*r*_ purely for visualisation (to spread out the long right tail); this display choice does not change the values of *Y* or affect any model estimates.

### 4.8 Statistical modeling and analysis

We summarize how the mean period varies with clustering using a central 80% (10th–90th percentile) contrast on *C*: from *C*_0.10_ to *C*_0.90_. On the natural-log scale this maps directly to a ratio on the period scale [27]. The regression is fitted to all graphs; for the *C*_0.10_–*C*_0.90_ contrast we use the fitted clustering slope *β*_*C*_ and the observed span (*C*_0.90_ − *C*_0.10_), define Δ*Y* = *β*_*C*_ (*C*_0.90_ − *C*_0.10_), and report exp(Δ*Y*) as the adjusted ratio on the period scale. In cross-validation we compute the same contrast separately in each fold using the fold-specific *β*_*C*_. Plots display the fitted curve across the full observed support of *C* and highlight the central 80% window used for the contrast.

#### Main regression for mean period

We model the average log-period *Y* (Eq. (9)) with a Gaussian additive model

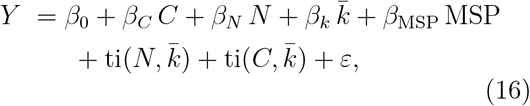

where *N* is the number of nodes, 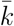 the average degree, and MSP the mean directed shortest path (Eq. (3)). We model the size-degree and clustering-degree interactions with tensor-product smooths ti(*N*, 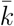) and ti(*C*, 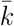) using cubic regression-spline marginals.

The basis dimensions for *N*, 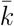, *C* and MSP are chosen adaptively based on the number of distinct observed values in each covariate and capped at moderate values (typically between 3 and 10 basis functions per margin) to allow mild nonlinearity without overfitting. Models are fitted by penalised least squares via restricted maximum likelihood (REML) with shrinkage selection, using the mgcv package in R with a Gaussian error distribution. Adjusted *C*_0.10_-*C*_0.90_ contrasts and confidence intervals are computed from *β*_*C*_ as described above and reported on the period scale as exp(Δ*Y*).

#### Robustness models

As robustness checks, we consider two alternative specifications. First, we replace the linear clustering term by a smooth *s*(*C*) while keeping the same interaction structure, to test whether a more flexible, nonlinear relationship between clustering and the log-mean period substantially improves the fit. Second, we fit a more flexible generalized additive model with separate smooth terms for *N*, 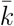, *C*, MSP and ABC and the same tensor-product interactions, treating this as a fully smooth benchmark. These variants are compared to the main model using both in-sample criteria and out-of-sample performance. To assess the importance of the interaction structure, we also compare the main model to reduced additive models without tensor-product terms, and to models without the ti(*C*, 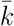) interaction, using likelihood-ratio *χ*^2^ tests (via `anova.gam in mgcv`) and report the corresponding test statistics and *p*-values for the interaction effects.

#### Unadjusted period summaries by clustering

Because the analysis dataset contains a single row per graph (the average natural log-period *Y* over *M* =100 runs; Sec. 4.7), we describe how period distributions change with clustering by grouping graphs into deciles of *C* and summarising the geometric mean period *L*_geo_ = exp(*Y*) within each decile. In particular, we report for each decile the number of graphs, the median, and the interquartile range of *L*_geo_ (Table 3). This gives a simple, nonparametric description of how typical periods and their spread vary across the observed range of *C*, without making extra assumptions about the shape of the distribution.

#### Validation and diagnostics

Model adequacy is assessed by 10-fold cross-validation at the graph level (folds contain disjoint sets of graphs), predicted-versus-observed comparisons, calibration by prediction percentiles, residual-versus-fitted plots, and normal Q–Q plots. In cross-validation we fit the main model with linear *C*, the smooth-*C* variant, and the fully smooth benchmark on the training folds and compare their out-of-sample performance using root mean squared error and *R*^2^ on the held-out graphs (graphs not used to fit the model). To check robustness to modeling choices, we use likelihood-ratio *χ*^2^ tests to compare the main specification to the smooth-*C* variant and to reduced models without interaction terms; these tests quantify the evidence for nonlinearity in *C* and for the significance of the interaction structure.

## 5 Results

We study how clustering coefficient variation in small-world graphs shapes the long-run dynamics of synchronous, signed threshold Boolean networks. Using the Gaussian additive model in Sec. 4 (Eq. (16)), we estimate the effect of clustering on the average log-period *Y* while adjusting for *N*, 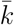, and mean directed shortest path (MSP), and allowing tensor product interactions ti(*N*, 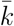) and ti(*C*, 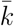) to capture size-degree and clustering-degree structure. Because changing 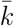 can drive both clustering and MSP in the WattsStrogatz model, we evaluate predictors jointly to address collinearity. Raw Spearman correlations between *Y* and each structural descriptor are all positive, ranging from *ρ* ≈ 0.48 for *N* and *ρ* ≈ 0.69 for MSP to *ρ* ≈ 0.94 for mean degree, with *ρ* ≈ 0.81 for *C* and *ρ* ≈ 0.85 for average betweenness. These strong positive correlations occur because larger, more highly connected graphs also tend to have higher clustering and shorter paths. For this reason we analyse all predictors together in one model, so that the effect of clustering is interpreted as a *partial* effect after adjusting for *N*, 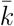 and MSP. Average betweenness centrality (ABC) had only a very small additional partial effect in exploratory fits once *N*, 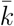, *C*, and MSP were included, so ABC is not retained in the main regression. Importantly, the positive raw association between *C* and *Y* reverses after adjustment: once *N*, 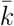, and MSP are held fixed, higher clustering is associated with *shorter* average periods. The remainder of this section first quantifies this adjusted clustering effect, then describes the roles of the other graph properties, model fit and diagnostics, and finally the unadjusted distributions of periods across clustering deciles.

### 5.1 Primary effect of clustering on attractor period

In the main model the regression term for clustering *C* is linear and negative, and we summarise this effect by the coefficient 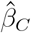. Numerically, the fitted coefficient is 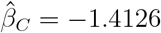 with standard error 0.6988, 95% confidence interval [−2.78, −0.04], and *p ≈* 0.044. A 0.10 increase in *C* multiplies the expected geometric mean attractor period *L*_geo_ = exp(*Y*) by

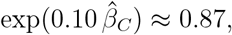

which corresponds to about a 13% reduction on the period scale. We also summarise the effect over the central 80% of the observed *C* values, from the 10th percentile *C*_0.10_ = 0.0000 to the 90th percentile *C*_0.90_ = 0.4599. Here *C*_0.10_ and *C*_0.90_ denote the 10th and 90th percentiles of the clustering coefficient across all graphs, so the contrast *C*_0.10_ → *C*_0.90_ compares a typical low-clustering graph to a typical high-clustering graph. Over this range the adjusted ratio on the period scale is

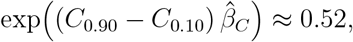

so moving from *C*_0.10_ to *C*_0.90_ shortens the expected geometric mean period by roughly 48%.

Grouped 10-fold cross-validation at the graph level gives very similar effect sizes: across folds the mean clustering slope is 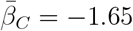, so a 0.10 increase in *C* multiplies the geometric mean period by exp 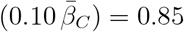 with 95% CI [0.76, 0.91], and the interdecile ratio is ≈ 0.49 with 95% CI [0.28, 0.65] (Table 2). Fold-specific slopes are negative in every split, ranging from *β*_*C*_≈ −0.93 to *β*_*C*_ ≈ −2.79, with the corresponding ratios exp(0.10 *β*_*C*_) lying between about 0.76 and 0.91 and the interdecile ratios between about 0.28 and 0.65, so the shortening effect is present across all folds rather than driven by a few particular graphs. Taken together, the model and cross-validated slopes correspond to about a 13-15% shortening in the geometric mean period per +0.10 increase in *C*.

The shape of the adjusted clustering effect is illustrated in Fig. 4, which plots the effect of *C* on predicted log-period after adjusting for *N*, 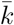 and MSP together with partial residuals for *C*. Each point shows how far a graph’s observed *Y* lies above or below the fitted model once other covariates are accounted for, while the curves show the estimated conditional effect of clustering from the linear-*C* model and from a more flexible smooth-*C* model. Both curves decline steadily with *C*, and the residual cloud follows the same downward trend, confirming that, conditional on the other graph properties, higher clustering is associated with shorter average log-periods. Additional diagnostics based on individual conditional expectation, accumulated local effects, and the numerical derivative of the smooth *s*(*C*) term show only mild curvature and a nearly constant negative slope over the observed *C* range; these robustness checks are described in Sec. 5.5.

### 5.2 Effects of graph properties

While the focus is clustering, the main Gaussian additive model (Eq. (16)) also quantifies how size *N*, mean degree 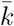, and mean directed shortest path (MSP; Eq. (3)) relate to periods when considered together. In this specification we include linear terms for *N*, 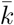, *C*, and MSP plus the tensor-product interactions ti(*N*, 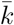) and ti(*C*, 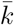). Table 1 reports the parametric coefficients for the linear terms.

**Table 1.** Parametric coefficients for the mean log-period model. Outcome *Y* is the average log-period as in Eq. (9); predictors: *N*, 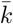, *C*, and MSP. Entries are adjusted per unit effects on the log scale with Wald 95% CIs; the last column reports the proportional change exp(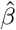) per +1 unit. For *C*, results interpret the prespecified +0.10 contrast via exp 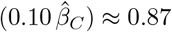.

| Term | Estimate | Std. Error | 95% CI low | 95% CI high | Ratio per +1 |
| --- | --- | --- | --- | --- | --- |
| Intercept | -2.754 | 0.150 | -3.049 | -2.459 | - |
| $N$ | 0.0173 | 0.00188 | 0.0136 | 0.0210 | 1.017 |
| $\bar{k}$ | 0.750 | 0.0552 | 0.642 | 0.859 | 2.12 |
| $C$ | -1.413 | 0.699 | -2.78 | -0.043 | 0.24 |
| MSP | 0.0555 | 0.0285 | -0.0004 | 0.111 | 1.06 |

In the linear plus tensor model, *N*, 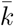, and MSP all have positive associations with *Y* (average log-period), whereas clustering has a negative association consistent with Sec. 5.1. The fitted coefficients are 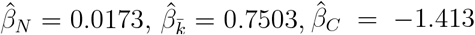, and 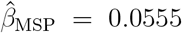 (Table 1). On the geometric-mean period scale, a +10 increase in *N* multiplies the expected period by exp 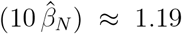, about a 19% increase, a +1 increase in mean degree multiplies it by 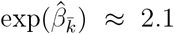, more than doubling the typical period, and a +1 increase in MSP multiplies it by 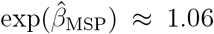, a modest effect compared with degree. For clustering, the per unit ratio 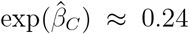 corresponds to the smaller +0.10 contrast discussed above, where 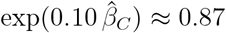, about a 13% reduction. As before, we do not compare raw per unit magnitudes across predictors because units differ; the proportional change column in Table 1 provides a compact way to read off effects on the period scale.

The interaction ti(*N*, 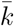) indicates that the influence of size depends on degree: when 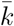 is near 2 the effect of increasing *N* is negligible, but at moderate to high 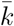 the effect of *N* becomes clearly positive, consistent with a positive size-degree interaction. The ti(*C*, 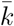) term captures that the influence of clustering depends on degree: at low 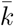 changing *C* has little impact, whereas at higher 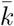 increasing clustering pulls the fitted surface toward shorter log-periods, reinforcing the negative conditional effect of *C* found in Sec. 5.1. These patterns are visualised in Fig. 1, which shows the predicted log(mean period) over (*C*, 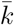) at fixed *N* and MSP, and in Fig. 2, which shows the corresponding surface over (*N*, 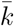) with the simulated design points overlaid. Dense, highly connected networks can support long attractor periods, but in this model adding triangles pulls the dynamics back toward shorter cycles. Partial-dependence profiles and tensor-product surfaces from the main linear-plus-tensor model show the same pattern: mean degree has the strongest effect, MSP and *N* have positive but milder effects, and once these are accounted for the additional contribution of average betweenness is small. To quantify the additional fit provided by the interaction terms, we also compare the main model to reduced additive models without tensor-product terms and to a model without the ti(*C*, 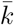) interaction using likelihood-ratio *χ*^2^ tests (`anova.gam` in R); the corresponding test statistics and *p*-values are reported in Appendix Table 4, which summarises the statistical significance of the interaction structure.

**Figure 1.**
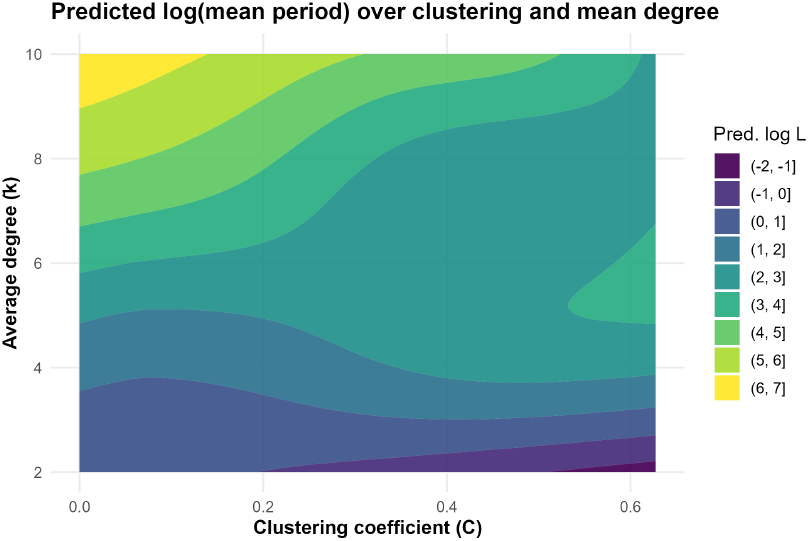
Predicted log(mean period) over clustering and mean degree. Contour surface of the fitted linear-plus-tensor model over clustering coefficient *C* (horizontal axis) and mean degree 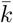 (vertical axis), with other covariates held at their medians. Periods increase strongly with degree, while at any fixed degree increasing *C* shifts the surface downward, especially at moderate and high 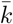, illustrating that higher clustering shortens the predicted average period in denser networks.

**Figure 2.**
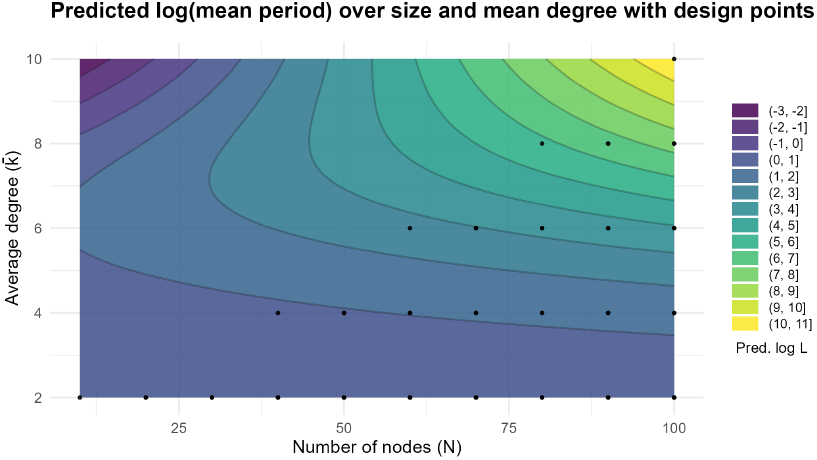
Predicted log(mean period) over size and mean degree with design points. Filled contours of the fitted linear-plus-tensor model over network size *N* (horizontal axis) and mean degree 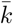 (vertical axis), with the other covariates held at their medians. Black points mark the simulated (*N*, 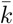) combinations. Predicted periods increase with both *N* and 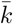, and the effect of size is strongest at higher mean degree. The design points show that much of the plotted surface is supported by simulated graphs; however, the upper-left region (small *N*, higher 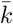) and a few boundary areas lack design points and should be viewed as extrapolation beyond the observed design.

### 5.3 Model fit

The additive Gaussian model for *Y* captures most of the between-graph variation in mean attractor period. For the primary linear plus tensor model (Eq. (16)) the in-sample performance is strong (adjusted *R*^2^ = 0.94, deviance explained 94.3%, RMSE on the log scale ≈ 0.51, scale estimate ≈ 0.27, *n* = 330). Standard predictedversus-observed comparisons, calibration by prediction percentiles, residual-versus-fitted plots, and normal Q-Q plots show good agreement and no major systematic deviations, supporting the adequacy of the Gaussian GAM on the log scale. These goodness-of-fit diagnostics (predicted vs. observed, calibration by deciles, residuals vs. fitted values, and a normal Q-Q plot) are shown in Appendix Figs. 5-8; together they indicate a robust fit with good calibration and approximately Gaussian residuals. When graphs are grouped into deciles of the prediction, mean observed and mean predicted log-period agree closely, with differences generally within about 0.07 log units and at most ≈ 0.16, indicating that calibration holds well across the spectrum of fitted values.

Grouped 10-fold cross-validation at the graph level shows that out-of-sample performance is similar to the in-sample fit and that the estimated clustering effect is stable across folds. Table 2 summarises the cross-validated slopes *β*_*C*_ and the implied period ratios for a +0.10 increase in *C* and for the interdecile contrast *C*_0.10_ → *C*_0.90_.

**Table 2.** Cross-validated clustering effect. Grouped 10-fold cross-validation at the graph level for the main linear-plus-tensor model. For each fold we refit the model, record the clustering slope *β*_*C*_, and derive the corresponding period ratios for a +0.10 increase in *C* and for the interdecile contrast *C*_0.10_ → *C*_0.90_. The table reports the mean across folds, the standard deviation (SD), and an empirical 95% interval (2.5th-97.5th percentiles).

| Quantity | Mean | SD | 2.5% | 97.5% |
| --- | --- | --- | --- | --- |
| $\beta_C$ | -1.65 | 0.52 | -2.61 | -0.79 |
| Ratio for +0.10 in $C$ | 0.85 | 0.04 | 0.76 | 0.91 |
| Ratio for $C_{0.10} \rightarrow C_{0.90}$ | 0.49 | 0.10 | 0.28 | 0.65 |

### 5.4 Attractor period distributions by clustering

As a descriptive complement to the regression model, we also look directly at the raw data. In this subsection we study the *unadjusted* distributions of the graph-level geometric mean attractor period *L*_geo_ = exp(*Y*) across different values of the clustering coefficient *C*.

We sort graphs by *C* and split them into ten ordered groups labelled `D01-D10` using a rank-based decile cut on *C* (graphs with exactly the same *C* value stay together). Because many graphs have *C* ≈ 0, only eight of these deciles contain graphs: `D02` is the first non-empty bin (*n* = 112), and the remaining non-empty bins are `D04-D10`, which mostly contain about 33 graphs each, except D04 which has 21 (counts and summaries are given in Table 3).

**Table 3.** Decile sizes and summaries of geometric mean period by clustering decile. *n* is the number of graphs; medians and interquartile ranges (IQRs) are for *L*_geo_; *C*_mid_ is the bin midpoint (3 s.f.).

| Decile | $n$ | Median $L_{\text{geo}}$ | IQR $L_{\text{geo}}$ | $C_{\text{mid}}$ |
| --- | --- | --- | --- | --- |
| D02 | 112 | 1.00 | 0.000 | 0.000 |
| D04 | 21 | 1.07 | 0.466 | 0.0239 |
| D05 | 32 | 2.54 | 2.92 | 0.0485 |
| D06 | 33 | 6.55 | 29.3 | 0.0795 |
| D07 | 33 | 6.91 | 60.0 | 0.130 |
| D08 | 33 | 7.62 | 26.5 | 0.241 |
| D09 | 33 | 16.1 | 22.9 | 0.388 |
| D10 | 33 | 37.3 | 66.2 | 0.530 |

Figure 3 shows violin and box plots of *L*_geo_ by clustering decile, using a log_10_ *y*-axis for display. As clustering increases, both the typical period and the spread grow. In the lowest non-empty decile (`D02`) almost all graphs have very short periods (median = 1.00, interquartile range = 0.000). In the highest decile (`D10`) the median rises to about 37.3 and the interquartile range is over 60 time steps, with a long right tail. The intermediate deciles move smoothly between these extremes. Table 3 summarises the same pattern numerically: median *L*_geo_ rises from 1.00 to ≈ 37.3, and the interquartile range grows from essentially zero to more than 60 time steps.

**Figure 3.**
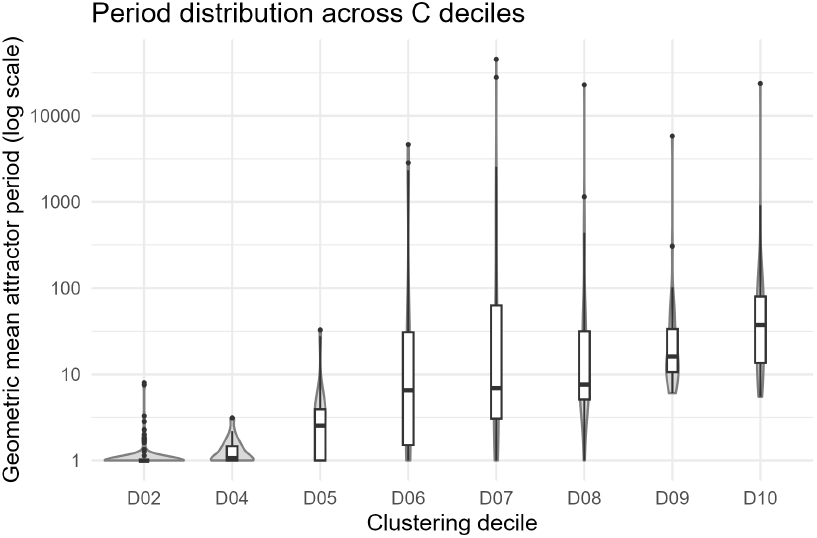
Geometric mean period by clustering decile (log_10_ *y*-axis). Violin and box plots of the geometric mean attractor period *L*_geo_ for each clustering decile. As clustering *C* increases, both the median and the spread of *L*_geo_ increase, so long typical periods are much more common in the high-clustering groups. The log_10_ *y*-axis is used only to spread out the long right tail for visual clarity and does not affect the underlying data or the regression model.

These raw patterns are descriptive, not causal. They show that, *without* adjustment, graphs with higher clustering tend to have longer and more variable periods, consistent with the positive Spearman correlation between *C* and *Y* reported earlier. In the Watts-Strogatz model, however, clustering does not change in isolation: higher *C* is typically accompanied by changes in network size, degree, and path structure. The adjusted analysis that conditions on *N*, 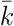, and MSP (Sec. 4) shows that the *partial* effect of clustering is in the opposite direction: when these other properties are held fixed, higher *C* shortens the expected geometric mean period (Sec. 5.1).

### 5.5 Robustness

We ask whether the estimated clustering effect could be an artefact of our modelling choices, in particular the interaction terms and the assumption of a linear effect in *C*.

First, we compare the main linear-plus-tensor model to simpler models using likelihood-ratio *χ*^2^ tests (Table 4). Adding the tensor-product terms ti(*N*, 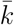) and ti(*C*, 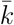) to an additive model with only linear effects of *N*, 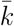, *C* and MSP reduces the deviance by about 125.5 units for 15.9 effective degrees of freedom (*p <* 0.001). Adding the ti(*C*, 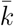) interaction on top of a model that already includes ti(*N*, 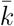) reduces deviance by a further 68.0 units for 12.8 degrees of freedom (*p <* 0.001). These tests support keeping both interactions and show that the effect of clustering depends on degree in a statistically important way.

Second, we then test whether a more flexible, nonlinear term in clustering improves the fit. We replace the linear *C* term by a smooth function *s*(*C*) in an otherwise identical Gaussian additive model (Sec. 4, Eq. (16)), keeping the same linear terms for *N*, 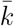 and MSP and the same tensor-product interactions ti(*N*, 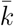) and ti(*C*, 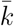). The smooth *s*(*C*) is a cubic spline with basis size *k*_*C*_, and it is penalised so that the fitted curve is kept as smooth as possible and does not introduce unnecessary wiggles. The fitted effective degrees of freedom for *s*(*C*) are only slightly above 1, which means the estimated effect of *C* is almost linear, with only mild curvature. Deviance explained and adjusted *R*^2^ are essentially unchanged compared with the linear-*C* model, and the smooth-*C* variant has a slightly *higher* residual deviance (Table 5; Δdeviance ≈ −0.14 at Δ*df* ≈ 0.29). There is therefore no evidence that a nonlinear effect in *C* improves the fit, and we retain the linear specification for *C*.

**Table 4.** Likelihood-ratio tests for interaction terms. Each row compares a reduced model with the main linear-plus-tensor model for *Y* (Eq. (16)). The first comparison tests the joint contribution of both interaction terms ti(*N*, 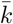) and ti(*C*, 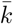). The second comparison tests the additional contribution of the clustering-degree interaction ti(*C*, 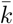) when ti(*N*, 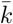) is already included. The LR statistic is the change in deviance (*χ*^2^) with approximate degrees of freedom (df).

| Comparison | Full model | Reduced model | Terms tested<br>(dropped) | $\chi^2$ | df | $p$-value |
| --- | --- | --- | --- | --- | --- | --- |
| Additive vs. full | Main model:<br>linear terms +<br>$\text{ti}(N, \bar{k})$ +<br>$\text{ti}(C, \bar{k})$ | Additive model:<br>linear terms only<br>(no ti terms) | $\text{ti}(N, \bar{k})$, $\text{ti}(C, \bar{k})$ | 125.5 | 15.9 | < 0.001 |
| No $\text{ti}(C, \bar{k})$ vs. full | Main model:<br>linear terms +<br>$\text{ti}(N, \bar{k})$ +<br>$\text{ti}(C, \bar{k})$ | Main model<br>without $\text{ti}(C, \bar{k})$ | $\text{ti}(C, \bar{k})$ | 68.0 | 12.8 | < 0.001 |

**Table 5.** Linear vs smooth clustering effect. Comparison of the main model with a linear term in clustering coefficient *C* and a variant with a penalised spline *s*(*C*). The smooth-*C* variant has slightly higher residual deviance, so the linear term is preferred.

| Model | Residual df | Residual deviance | $\Delta\text{df}$ | $\Delta\text{deviance}$ |
| --- | --- | --- | --- | --- |
| Linear $C$ | 309.09 | 84.21 | - | - |
| Smooth $s(C)$ | 308.80 | 84.36 | 0.29 | -0.14 |

The shape of the fitted clustering effect is also consistent with this conclusion. Figure 4 plots partial residuals for *C* from the linear model together with the estimated contributions of the linear *C* term and the smooth *s*(*C*) term. Both fitted curves decline steadily with *C*, they are almost straight over the observed range, and the residual cloud follows the same downward trend. This indicates an approximately constant negative slope on the log scale as clustering increases, and shows that the linear term in *C* is a parsimonious and adequate summary of the conditional effect.

**Figure 4.**
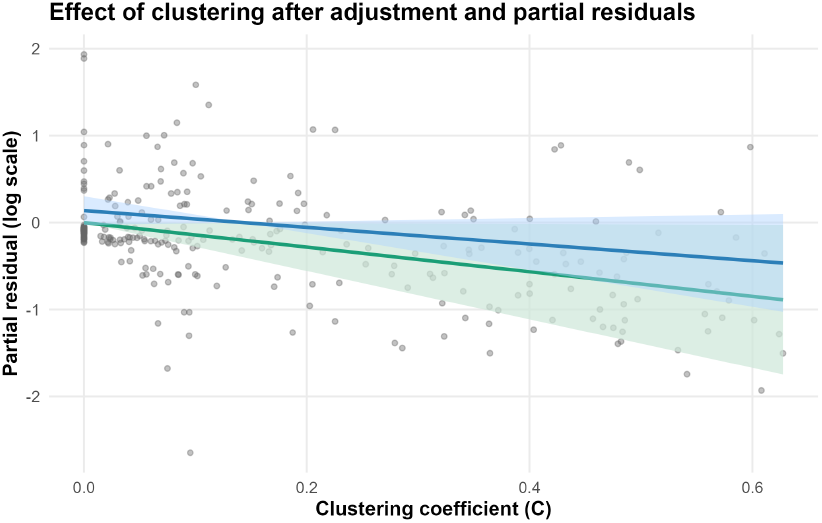
Effect of clustering after adjustment and partial residuals. Partial residuals for *C* from the linear model (grey points), plotted against *C*, together with the estimated conditional effect of clustering from the linear-*C* model (green curve) and from the smooth-*C* model (blue curve), each with a 95% confidence band. The residual cloud follows a clear downward trend, and both fitted curves decline steadily with *C*, confirming a negative conditional association between clustering and the average log-period after adjusting for *N*, 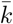 and MSP.

Finally, we fitted a more flexible comparison model with separate smooth terms for *N*, 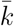, *C*, MSP and ABC, plus the same tensor-product interactions. This fully smooth model gives similar deviance explained and out-of-sample performance to the main linear-plus-tensor specification. Partial-dependence profiles for the additional covariates (Appendix Fig. 9) show that mean degree has the strongest effect on *Y*, MSP and *N* have positive but milder effects, the partial effect of ABC is small once the other descriptors are included, and the negative effect of *C* remains, consistent with the main model.

#### Reproducibility

All code and data will be publicly available. A tagged GitHub release will include scripts to rebuild all tables and figures, the processed data, a manifest (commit hash and random seeds), and an environment file with package versions. We will archive the release with a DOI and cite it; until the DOI is issued, materials are available on request.

## 6 Discussion

We quantify how small-world wiring controls cycle length in synchronous, signed threshold Boolean networks. Adjusting for size *N*, mean degree 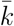, and mean directed shortest path (MSP) in a single model with size-degree and clustering-degree tensor terms (Sec. 4), higher clustering *C* is linked to shorter attractor periods. In the main regression, a 0.10 step in *C* multiplies the expected geometric mean period *L*_geo_ = exp(*Y*) by exp 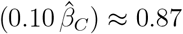 (about 13% shorter), and moving from *C*_0.10_ = 0.0000 to *C*_0.90_ = 0.4599 yields an average ≈ 48% reduction (exp(Δ*Y*) ≈ 0.52; Sec. 5). Cross-validated estimates are very similar: grouped 10-fold CV at the graph level gives a mean slope 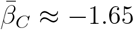, so a 0.10 increase in *C* multiplies the geometric mean period by exp 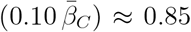 (Table 2). The shortening effect of clustering therefore generalises across held-out graphs and corresponds to about a 13-15% shortening in the geometric mean period per +0.10 increase in *C*.

Because the WS rewiring probability alters triangle density and path length together, *C* and MSP co-vary. We therefore identify the effect of clustering by modelling the *realised* graph statistics and estimating the *partial* association of *C* with the average log-period *Y* while conditioning on MSP (and on *N* and 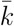). The negative slope for *C* persists under this joint adjustment, under cross-validation, and when the linear *C* term is replaced by a penalised smooth *s*(*C*). The smooth adds only slightly more than one effective degree of freedom and does not materially change the fitted effect or improve fit in terms of deviance explained, *R*^2^, or information criteria (Appendix Table 5). In Fig. 4 the fitted linear and smooth curves are almost indistinguishable, reinforcing that a linear term in *C* is adequate. We therefore interpret 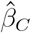 as a robust *conditional association* between clustering and shorter periods within the WS model.

Mechanistically, the pattern aligns with the ordered-chaotic spectrum for Boolean dynamics [3, 9, 2, 28]. Triangles create short feedback loops and local redundancy; together these features lower effective sensitivity, slow damage spreading, and shorten transients [25, 26, 35]. Empirically, the fitted curves for *C* in Fig. 4 are close to linear and remain negative over the observed range, suggesting that within our WS range clustering acts as a steady, order-promoting factor rather than a sharp switch. This complements prior work on information storage and perturbation spreading by providing a calibrated effect size on cycle length while controlling for size, degree, and path structure.

Other structural levers behave coherently. Holding the remaining predictors at their medians, increasing 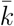 by +1 multiplies the expected geometric mean period by exp(0.750) ≈ 2.12, adding +10 nodes multiplies it by exp(10 × 0.0173) ≈ 1.19, and MSP is positively associated (ratio exp(0.0555)≈1.06 per unit) (Table 1). The fitted interaction between *N* and 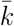 indicates a positive size-degree interaction: increases in *N* matter most at moderate to high 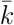. The corresponding clustering-degree interaction shows that the influence of clustering depends on degree: at low 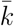 varying *C* has little impact, whereas at higher 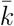 increasing *C* moves predictions toward shorter log-periods, consistent with a negative conditional effect of clustering.

Our study has a specific scope: we work with directed, signed WS graphs with synchronous updates and strict threshold rules over the ranges reported in Sec. 4. Model fit and calibration are strong in grouped 10-fold CV at the graph level (Sec. 5.3, Table 2), but extrapolations to asynchronous updates [29], other logic families (e.g. canalising or nested-canalising) [21, 37], heavy-tailed degree sequences [32, 14, 6], or strong modularity [33] should be made with care.

In sum, within the WS model analysed, clustering exerts a clear, independently estimable, monotone shortening effect on attractor periods once size, degree, and path length are accounted for. This links a concrete structural motif, triangles, to a quantitative, predictive handle on long-run dynamics in threshold networks relevant to synthetic circuits and interpretable models of cellular decision making [19, 5, 38].

## 7 Conclusion

Clustering is a simple, interpretable wiring feature with a clear dynamical consequence in signed, threshold Boolean systems. In Watts-Strogatz small-world graphs with synchronous updates, the *partial* effect of global clustering on the average log-period *Y* is monotone and close to linear, remains negative over the observed support (0 ≤ *C* ≤ 0.6274), and persists after conditioning on *N*, 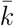, and MSP. These findings are consistent with theory on information storage, damage spreading, and loop closure. Because the effect is expressed in units of *C* itself, it is portable to empirical networks where clustering is measured directly. Practically, clustering joins degree and size, as well as path-length structure captured by MSP, as a quantitative control for tuning convergence speed and the length of repeating cycles in discrete dynamical systems.

## Acknowledgements

Maram Alqarni is supported by a scholarship from the Saudi Arabian Cultural Mission (SACM). This research was supported by the Australian Research Council through the Australian Research Council Centre of Excellence for Plant Success in Nature and Agriculture (CE200100015).

## Appendix

### A Model comparisons and robustness

#### A.1 Likelihood-ratio tests for interaction terms

Here we summarise the likelihood-ratio (LR) *χ*^2^ tests that compare the main linear-plus-tensor model with simpler models. The LR statistic is the change in deviance (*χ*^2^) with approximate degrees of freedom (df); *p*-values are from likelihood-ratio *χ*^2^ tests (using `anova.gam` in R).

The first comparison shows that adding both interaction terms ti(*N*, 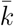) and ti(*C*, 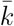) gives a large reduction in deviance (*χ*^2^ ≈ 125.5 for about 15.9 degrees of freedom, *p <* 0.001) compared with an additive model with only linear effects of *N*, 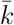, *C*, and MSP. The second comparison shows that adding ti(*C*, 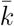) on top of ti(*N*, 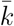) alone also gives a strong improvement in fit (*χ*^2^ ≈ 68.0 for about 12.8 degrees of freedom, *p <* 0.001). These tests support the use of the interaction structure in the main model.

#### A.2 Linear vs smooth clustering effect

We also compared the main model with a linear term in the clustering coefficient *C* to a variant where *C* is replaced by a penalised spline *s*(*C*). The smooth-*C* model has a slightly larger residual deviance and almost the same degrees of freedom, so there is no evidence that allowing extra curvature in *C* improves the fit.

The change in deviance is negative (Δdeviance ≈ −0.14 for a small change in df), so the smooth-*C* model does not give a better fit. This supports keeping a simple linear term in *C* in the main specification.

#### A.3 Model diagnostics

In this section we show the main goodness-of-fit diagnostics for the Gaussian additive model for *Y* (average log-period). All logarithms here are natural logs (log base *e*).

#### A.4 Predicted vs observed and calibration

Figure 5 shows the predicted versus observed log-period for all graphs, with a simple linear trend line. Points lie close to the diagonal, indicating a good overall fit. Figure 6 shows calibration by prediction deciles: graphs are grouped into ten bins by their predicted value, and we plot mean observed versus mean predicted log-period in each bin.

**Figure 5.**
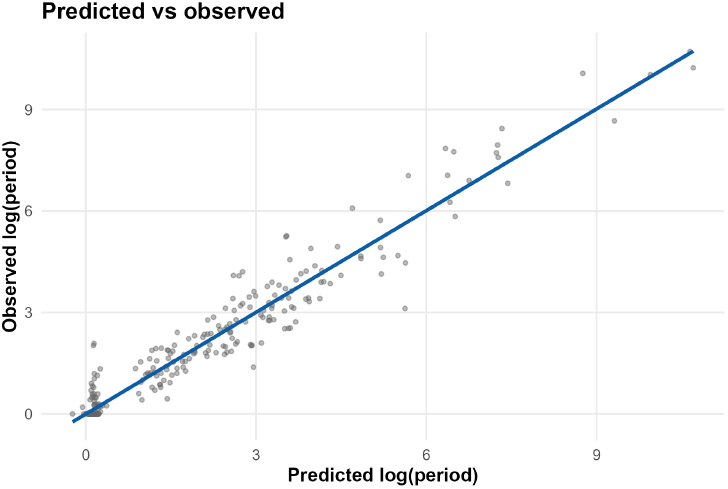
Predicted vs observed average log-period. Scatter plot of predicted versus observed *Y* = log(period) for all graphs under the main linear-plus-tensor model, with a fitted straight line. Points lie close to the diagonal, indicating good overall fit.

**Figure 6.**
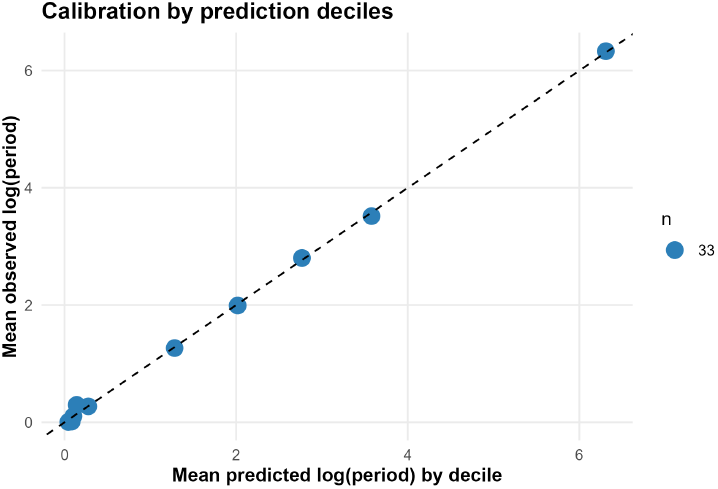
Calibration by prediction deciles. Mean observed versus mean predicted log-period by deciles of the predicted value. Point size shows the number of graphs in each decile. Most points lie close to the diagonal, and differences are small across the range, indicating good calibration.

#### A.5 Residual diagnostics

Figure 7 plots deviance residuals against the linear predictor. There is no strong pattern and the smooth trend line is close to zero across the range, suggesting that the mean structure is adequate. Figure 8 shows a normal Q-Q plot of the residuals. Points stay close to the reference line, except for mild deviations in the far tails, so the Gaussian assumption on the log scale is reasonable.

**Figure 7.**
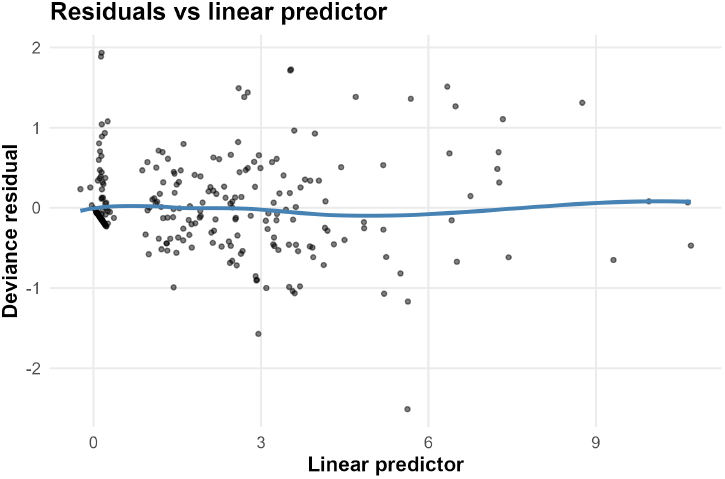
Residuals vs linear predictor. Deviance residuals of the main model versus the linear predictor. The smooth curve is close to zero and there is no strong structure, indicating that the model captures the main trends in *Y*.

**Figure 8.**
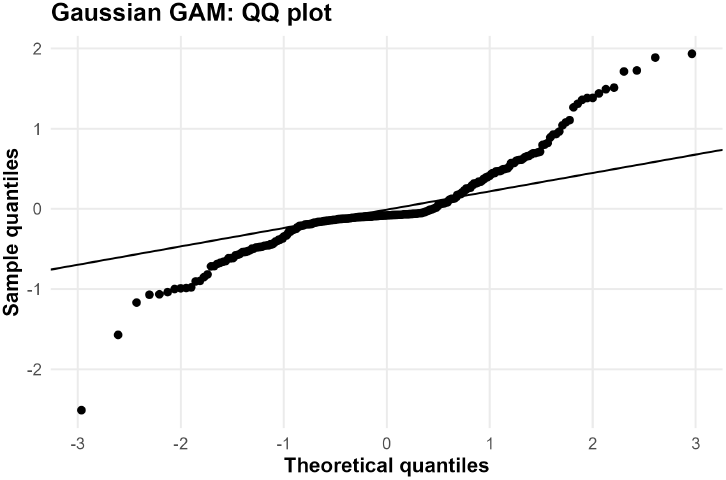
Normal Q-Q plot of deviance residuals. Normal Q-Q plot for the deviance residuals of the main model. Points are close to the reference line, with mild deviations in the tails, consistent with (though not definitive evidence for) a Gaussian error model on the log scale.

#### A.6 Additional partial-dependence plot

Figure 9 shows partial-dependence profiles for *N*, 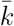, MSP, and average betweenness centrality (ABC), holding the other covariates at their median values. These plots illustrate that mean degree has the strongest effect on *Y*, MSP and *N* have positive but milder effects, and the additional contribution of ABC is small once the other variables are included.

**Figure 9.**
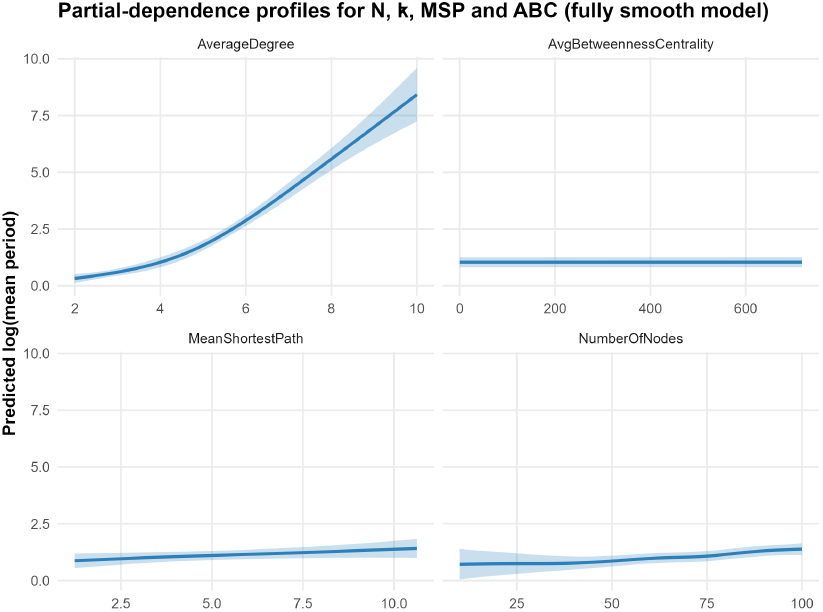
Partial-dependence profiles for graph descriptors. Partial-dependence curves for *Y* = log(period) as a function of each covariate (*N*, 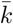, MSP, and ABC), with other covariates fixed at their medians. Shaded bands are 95% confidence intervals. Mean degree 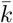 has the strongest effect, MSP and *N* are positive but milder, and the effect of ABC is small once the other descriptors are included.

## References

[1] Carlos Gershenson. Introduction to random boolean networks. arXiv preprint nlin/0408006, 2004.

[2] Barbara Drossel. Random boolean networks. Reviews of nonlinear dynamics and complexity, pages 69–110, 2008.

[3] S. A. Kauffman. Metabolic stability and epigenesis in randomly constructed genetic nets. Journal of Theoretical Biology, 22(3): 437–467, 1969.

[4] Martin Hopfensitz, Christoph Müssel, Markus Maucher, and Hans A Kestler. Attractors in boolean networks: a tutorial. Computational Statistics, 28:19–36, 2013.

[5] Assieh Saadatpour and Réka Albert. Boolean modeling of biological regulatory networks: a methodology tutorial. Methods, 62(1):3–12, 2013.

[6] Maximino Aldana. Boolean dynamics of networks with scale-free topology. Physica D: Nonlinear Phenomena, 185(1):45–66, 2003.

[7] Konstantin Klemm and Stefan Bornholdt. Stable and unstable attractors in boolean networks. Physical Review E, 72(5):055101, 2005.

[8] Phillip Bonacich. Some unique properties of eigenvector centrality. Social networks, 29 (4):555–564, 2007.

[9] Ilya Shmulevich and Stuart A Kauffman. Activities and sensitivities in boolean network models. Physical review letters, 93(4):048701, 2004.

[10] Stefan Bornholdt and Kim Sneppen. Robustness as an evolutionary principle. Proceedings of the Royal Society of London. Series B: Biological Sciences, 267(1459):2281–2286, 2000.

[11] Fangting Li, Tao Long, Ying Lu, Qi Ouyang, and Chao Tang. The yeast cell-cycle network is robustly designed. Proceedings of the National Academy of Sciences, 101(14): 4781–4786, 2004.

[12] Reinhard Diestel. Graph Theory. Springer, 5th edition, 2017.

[13] Frank Harary. Graph Theory. Addison-Wesley, 1969.

[14] Réka Albert and Albert-László Barabási. Statistical mechanics of complex networks. Reviews of modern physics, 74(1):47, 2002.

[15] Mark EJ Newman. The structure and function of complex networks. SIAM review, 45 (2):167–256, 2003.

[16] Mikaela Koutrouli, Evangelos Karatzas, David Paez-Espino, and Georgios A Pavlopoulos. A guide to conquer the biological network era using graph theory. Frontiers in bioengineering and biotechnology, 8:34, 2020.

[17] Duncan J Watts and Steven H Strogatz. Collective dynamics of ‘small-world’ networks. Nature, 393(6684):440–442, 1998.

[18] Mark EJ Newman. The mathematics of networks. The new palgrave encyclopedia of economics, 2(2008):1–12, 2008.

[19] Leon Glass and Stuart A Kauffman. The logical analysis of continuous, non-linear biochemical control networks. Journal of theoretical Biology, 39(1):103–129, 1973.

[20] S. Karanam and W.-J. Rappel. Boolean modelling in plant biology. Journal of Theoretical Biology, 2022.

[21] Stuart Kauffman, Carsten Peterson, Björn Samuelsson, and Carl Troein. Genetic networks with canalyzing boolean rules are always stable. Proceedings of the National Academy of Sciences, 101(49):17102–17107, 2004.

[22] Tiago P Peixoto and Barbara Drossel. Boolean networks with reliable dynamics. Physical Review E—Statistical, Nonlinear, and Soft Matter Physics, 80(5):056102, 2009.

[23] Hadeel Kittaneh, Filippo Castiglione, and Abdul Salam Jarrah. Stability of threshold boolean networks. Journal of Complex Networks, 13(4):cnaf011, 2025.

[24] Xiao Yang, Nilam Ram, Peter CM Molenaar, and Pamela M Cole. Describing and controlling multivariate nonlinear dynamics: A boolean network approach. Multivariate Behavioral Research, 57(5):804–824, 2022.

[25] Joseph T Lizier, Siddharth Pritam, and Mikhail Prokopenko. Information dynamics in small-world boolean networks. Artificial life, 17(4):293–314, 2011.

[26] Qiming Lu and Christof Teuscher. Damage spreading in spatial and small-world random boolean networks. Physical Review E, 89(2): 022806, 2014.

[27] Gareth James, Daniela Witten, Trevor Hastie, and Robert Tibshirani. An introduction to statistical learning: with applications in R, volume 103. Springer, 2013.

[28] Florian Greil and Kevin E Bassler. Attractor period distribution for critical boolean networks. Physical Review Letters, 102(4): 048101, 2009.

[29] Albert I. Albert R. Saadatpour, A. Attractor analysis of asynchronous boolean models of signal transduction networks. Journal of Theoretical Biology, 266(4):641–656, 2010.

[30] Deok-Sun Lee. Evolution of regulatory networks towards adaptability and stability in a changing environment. Physical Review E, 90(5):052822, 2014.

[31] Jason Lloyd-Price, Abhishekh Gupta, and Andre S Ribeiro. Robustness and information propagation in attractors of random boolean networks. PloS one, 7:1–6, 2012.

[32] Albert-László Barabási and Réka Albert. Emergence of scaling in random networks. Science, 286(5439):509–512, 1999.

[33] Neeraj Pradhan, Subinay Dasgupta, and Sitabhra Sinha. Modular organization enhances the robustness of attractor network dynamics. Europhysics Letters, 94(3):38004, 2011.

[34] Caitlyn Parmelee, Samantha Moore, Katherine Morrison, and Carina Curto. Core motifs predict dynamic attractors in combinatorial threshold-linear networks. PloS one, 17(3): e0264456, 2022.

[35] Carlos Handrey A Ferraz and Hans J Herrmann. The kauffman model on small-world topology. Physica A: Statistical Mechanics and its Applications, 373:770–776, 2007.

[36] Ugo Bastolla and Giorgio Parisi. Closing probabilities in the kauffman model: An annealed computation. Physica D: Nonlinear Phenomena, 98(1):1–25, 1996.

[37] Maximino Aldana and Philippe Cluzel. A natural class of robust networks. Proceedings of the National Academy of Sciences, 100 (15):8710–8714, 2003.

[38] Jorge GT Zañudo and Réka Albert. Cell fate reprogramming by control of intracellular network dynamics. PLoS computational biology, 11(4):e1004193, 2015.

[39] Andy Beatty, Christopher R Winkler, Thomas Hagen, and Mark Cooper. Predicting trait phenotypes from knowledge of the topology of gene networks. bioRxiv, page 2021.06.29.450449, 2021.

[40] Linton C Freeman. A set of measures of centrality based on betweenness. Sociometry, pages 35–41, 1977.

[41] Andrew Pomerance, Edward Ott, Michelle Girvan, and Wolfgang Losert. The effect of network topology on the stability of discrete state models of genetic control. Proceedings of the National Academy of Sciences, 106 (20):8209–8214, 2009.

